# A photostable version of HY5 confers tolerance to proximity shade and improved defense responses in tomato

**DOI:** 10.64898/2026.09.25.754491

**Authors:** Esteban Burbano-Erazo, Jamila Dich, Margalida Vidal-Tur, Miguel Simon-Moya, Antonella Anna Maria Locascio, Silvana Francesca, Maria Manuela Rigano, Mohamed Faize, Jaume F. Martinez-Garcia, Manuel Rodriguez-Concepcion

## Abstract

Light is essential for plant growth and development. Sustainably feeding a constantly-growing human population will likely involve adapting crop plants to intercropping and high planting density by rational manipulation of light signaling. Here, we edited the tomato (*Solanum lycopersicum*) genome to generate lines with a light-stable version of ELONGATED HYPOCOTYL 5 (HY5), a master transcription factor involved in the integration of light and hormone signaling. Removing the tomato HY5 N-terminal domain required for interaction with CONSTITUTIVE PHOTOMORPHOGENIC 1 (COP1) prevented light-dependent protein degradation and resulted in a gain-of-function phenotype of short seedlings. Elongation growth was also compromised under proximity shade conditions either simulated by enriching white light (W) with far-red light (W+FR) or achieved by growing plants at a higher density. Transcriptomic analysis of gene expression changes after exposure to W+FR for 24h revealed a reduced number of shade-responsive genes in edited lines compared to unedited, wild-type controls, many of which are related to growth and hormone (notably auxin) biosynthesis and signaling. The reduced elongation observed in edited lines correlated with enhanced resistance to infection by viral, bacterial and fungal pathogens, both under low and high density conditions. These results indicate that our editing approach allows the generation of gain-of-function tomato plants in which HY5 is camouflaged to avoid COP1 recognition and eventual degradation. Our findings therefore provide a biotechnological tool to create more compact and pathogen-resistant plants amenable to high planting densities.

## INTRODUCTION

Emerging agricultural systems, including high-density planting, intercropping, and vertical farming, involve complex light environments that differ from those found by crop plants under conventional cultivation (Chen et al., 2026). Altered light quantity and quality conditions derived from the proximity to neighboring plants promote elongation growth at the expense of reducing defense capacity through a growth–defense trade-off, resulting in an increased risk of infections associated with crowded (high planting density) environments (Pierik and Ballare, 2021). Understanding how plants sense and respond to changes in light conditions through signaling pathways is essential for developing new crop varieties adapted to growing closer together or under the canopy of other plants.

Plants perceive their surrounding light conditions using different classes of photoreceptors, including phytochromes that perceive and transduce red and far-red light (R and FR, respectively). Under dense vegetation, preferential absorption of R and reflection of FR by photosynthetic tissues results in a reduction in the R to FR ratio (R:FR) that is perceived by phytochromes. This low R:FR signal triggers a developmental program characterized by enhanced stem and petiole elongation, reduced branching, leaf hyponasty, and changes in resource allocation that also affect stress resilience and yield (Ballare and Pierik, 2017; Martinez-Garcia and Rodriguez-Concepcion, 2023). Light signals sensed and transduced by photoreceptors converge onto transcription factors such as ELONGATED HYPOCOTYL 5 (HY5), eventually modulating gene expression. In *Arabidopsis thaliana*, HY5 regulates a wide range of processes, including photomorphogenesis and responses to changing light and temperature cues (Ang et al., 1998; Lee et al., 2007; Gangappa and Botto, 2016; Ortiz-Alcaide et al., 2019; Burko et al., 2020; Xiao et al., 2022). Importantly, HY5 levels and activity are modulated by physical interactions with other proteins (Gangappa and Botto, 2016; Xiao et al., 2022).

The C-terminal bZIP (basic region/leucine zipper) domain of HY5 contains a basic region for DNA binding and a leucine zipper motif for protein-protein interaction that allows to form a functional homodimer. The N-terminal region contains a casein kinase II phosphorylation domain (ESDEE, residues 35–39 in the Arabidopsis protein) and a motif required for interaction with the E3 ubiquitin ligase CONSTITUTIVE PHOTOMORPHOGENIC 1 (COP1) that includes a valine–proline pair core (VP, residues 43–44) (Ang et al., 1998; Hardtke et al., 2000; Osterlund et al., 2000; Holm et al., 2001; Paulišić et al., 2021). HY5 physically interacts with COP1 and SPA1 through its N-terminal region, which contains for COP1 interaction that includes a valine–proline pair core (VP, residues 43–44) (Ang et al., 1998; Hardtke et al., 2000; Osterlund et al., 2000; Holm et al., 2001; Paulišić et al., 2021). COP1 binding in dark-germinated etiolated seedlings leads to HY5 ubiquitination and eventual 26S proteasome-mediated degradation to repress photomorphogenesis (Ang et al., 1998; Osterlund et al., 2000). COP1-mediated HY5 degradation is enhanced by binding to SUPPRESSOR OF PHYA-105 (SPA) proteins, a group of COP1 co-regulators (Saijo et al., 2003; Zhu et al., 2008). Upon illumination, the activity of the COP1/SPA complex is largely suppressed by photoreceptors and HY5 accumulates to induce photomorphogenesis, i.e., to repress Arabidopsis hypocotyl elongation and promote photosynthetic development (Podolec and Ulm, 2018). The S residue in the ESDEE motif is phosphorylated by SPA1, making HY5 relatively more stable in the dark (Hardtke et al., 2000; Wang et al., 2021). However, the unphosphorylated form of HY5 is the active physiological form, showing higher affinity for target promoters and to the COP1/SPA complex (Hardtke et al., 2000). The phosphorylated form of HY5 might serve as a reservoir pool in etiolated seedlings to be dephosphorylated quickly upon illumination for the initiation of seedling photomorphogenesis. Mutant HY5 proteins with the S36A substitution (i.e., constitutively unphosphorylated) are more stable and reduce the length of *hy5* hypocotyls beyond wild-type levels (Hardtke et al., 2000; Wang et al., 2021). Mutation of the VP core to AA also results in enhanced HY5 stability (Holm et al., 2001). While increased abundance of HY5 hardly impacts hypocotyl length in dark-grown Arabidopsis seedlings (Shi et al., 2018; Burko et al., 2020), HY5 represses hypocotyl elongation in fully de-etiolated seedlings exposed to low R:FR to simulate the presence of nearby vegetation (Ortiz-Alcaide et al., 2019; Morelli et al., 2021; Martinez-Garcia and Rodriguez-Concepcion, 2023). Interestingly, lines overexpressing HY5 resemble shade-tolerant plant species such as *Cardamine hirsuta* (Morelli et al., 2021).

Tomato (*Solanum lycopersicum*) is a popular crop worldwide that has emerged as an excellent model to investigate the molecular basis of proximity shade responses (Kunta et al., 2026; Mejia et al., 2026; Burbano-Erazo et al., 2025; Diaz-Lopez et al., 2026). As in Arabidopsis, tomato HY5 (SlHY5) is stabilized by light to regulate a variety of processes, including the response to proximity shade (Shiose et al., 2024; Yang et al., 2023; Zhang et al., 2022; Wang et al., 2018). Also like Arabidopsis, direct interaction with COP1 proteins results in SlHY5 degradation (Menconi et al.,2025). Here we investigated the feasibility of editing the tomato gene encoding SlHY5 to remove the N-terminal COP1-binding domain and hence generate a hyperstable yet functional transcription factor. The rationale behind this strategy was that preventing SlHY5 degradation in response to vegetation shade (low R:FR) would result in plants with enhanced tolerance to shade, which perform better when grown in close proximity to other plants (Burbano-Erazo et al., 2025).

## RESULTS

### Tomato lines with a genomic *SlHY5* variant lacking the N-terminal domain are shorter

Using CRISPR-Cas9, we edited the tomato gene encoding HY5 (*SlHY5*, Solyc08g061130) to create versions lacking the N-terminal domain of the protein required for interaction with the COP1/SPA complex. A single guide RNA (sgRNA) was designed targeting the *SlHY5* sequence encoding the predicted COP1 binding domain (**Figure 1A**). The same construct was used to transform the model tomato cultivar MicroTom (MT) and two commercial varieties, MoneyMaker (MM) and San Marzano (SM). Alleles harboring short insertions or deletions within the targeted sequence were identified and selected on the basis of their predicted protein products (**Supplemental Table S1**). The selected mutations shifted the reading frame, expectedly encoding protein variants in which the wild-type N-terminal domain was substituted by non-native sequences of different lengths resulting from the use of alternative translation start codons (**Figure 1A** and **Supplemental Table S1**). Homozygous edited lines lacking the Cas9 transgene were selected in MT (*hy5-mt1*, *hy5-mt3* and *hy5-mt4*), MM (*hy5-mm1* and *hy5-mm2*) and SM (*hy5-sm1* and *hy5-sm2*) backgrounds, and used for further experiments (**Figure 1A**).

**Figure 1.**
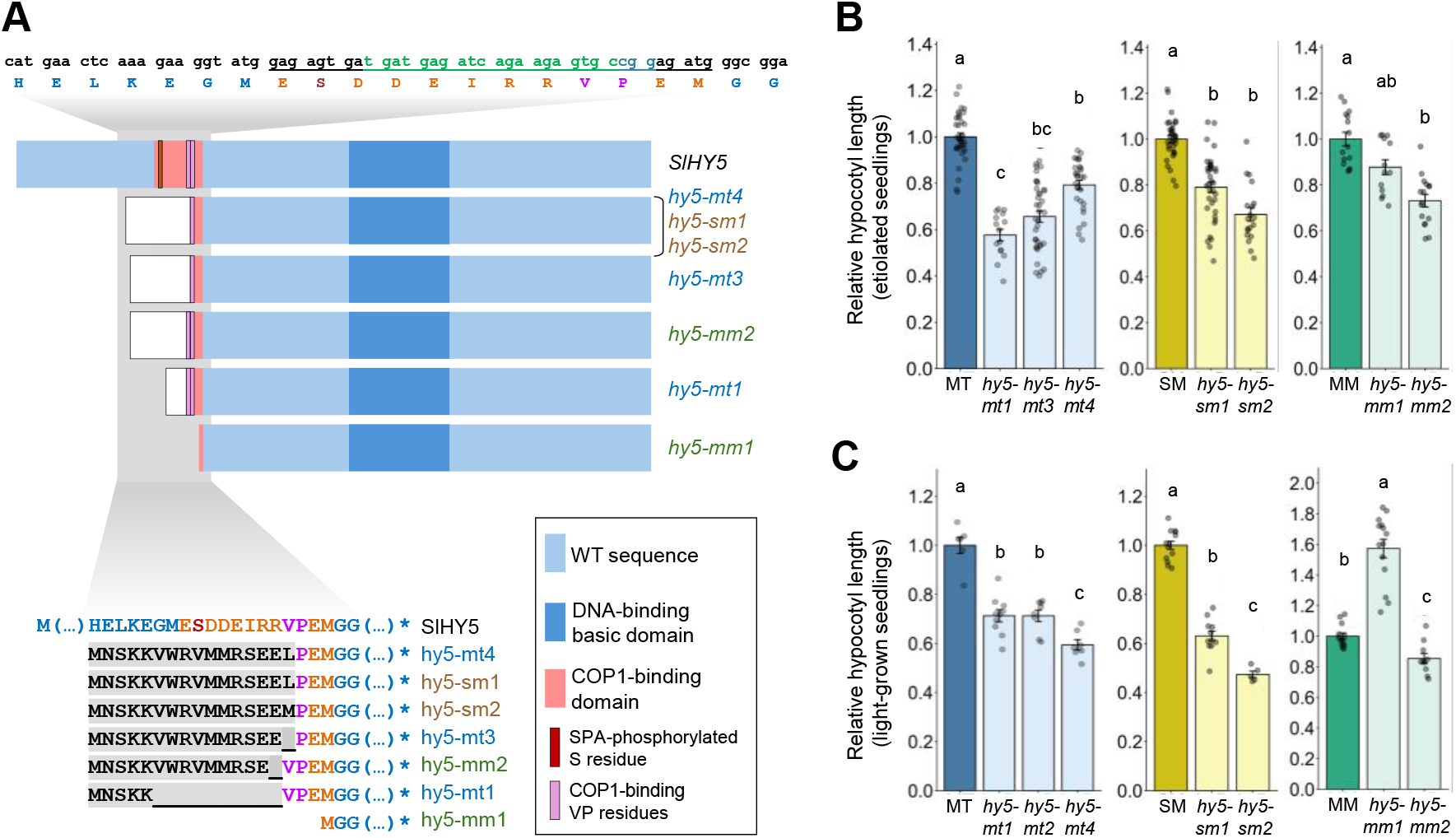
Tomato edited lines show altered elongation responses. (A) Schematic presentation of cDNA sequences from tomato wild-type (HY5) and edited lines (see also Table S1). The sgRNA target sequence is marked in green on the wild-type cDNA sequence shown on top. (B) Hypocotyl length of tomato seedlings germinated and grown in the dark for 5 days. (C) Hypocotyl length of tomato seedlings germinated and grown under long-day conditions for 11 days. In (B) and (C), hypocotyl length is represented relative to the corresponding WT parental (MicroTom, MT; San Marzano, SM; or MoneyMaker, MM), dots represent individual data points, error bars correspond to SD, and statistically significant difeerences are represented with leteers (one−way ANVAA followed by Duncanss multiple range test, *P* < 0.05).

To screen for lines with expectedly increased HY5 activity, we germinated and grew the selected mutants in darkness (**Figure 1B**). The rationale of this strategy was that a version of SlHY5 unable to interact with COP1 would not be degraded in the dark and hence it might accumulate in the nucleus to repress hypocotyl growth of etiolated seedlings. Under these conditions, all edited lines but *hy5-mm1* exhibited significantly shorter hypocotyls compared to their unedited parental line (**Figure 1B**). Notably, the mutation in the *hy5-mm1* line is the only one among the selected alleles that does not include a non-native N-terminal extension (**Figure 1A** and **Supplemental Table S1**). When the same lines were germinated and grown under long-day conditions (16h of light and 8h of darkness), *hy5-mm1* seedlings showed substantially longer hypocotyls than MM, whereas the rest of the alleles were shorter than their corresponding parental lines (**Figure 1C**). These results suggested that *hy5-mm1* might be a loss-of-function allele, in contrast to the putative gain-of-function phenotype of the lines carrying alleles with a predicted non-native N-terminal extension.

### Loss of interaction with tomato COP1 and SPA1 proteins results in SlHY5 photostability

To investigate whether the observed long and short hypocotyl phenotypes of CRISPR alleles resulted from differential protein accumulation, we compared the light-dependent stability of the proteins encoded by the two alleles available in the MM background (*hy5-mm1* and *hy5-mm2*). Multiple attempts using commercially-available antibodies against HY5 failed to detect the protein in tomato extracts. As an alternative, we transiently expressed the encoded proteins in *Nicotiana benthamiana* leaves (**Figure 2**). The cDNA sequences were isolated from MM, *hy5-mm1* and *hy5-mm2* seedlings (**Figure 2A**) and cloned with a myc tag fused to the C-terminus under the control of the *35S* promoter. The three generated constructs were agroinfiltrated in separate spots of several *N. benthamiana* leaves. After agroinfiltration, plants were left to recover for one day under long day conditions and then transferred to continuous white light (W) for two additional days. Just before the transfer, some of the agroinfiltrated leaves were covered with aluminum foil to remain in complete darkness while attached to the W-exposed plants. Samples from leaves grown in the light or in the dark were then collected and used for RNA and protein extraction. Transcript levels were similar for the three constructs in the two conditions tested (**Figure 2B**). By contrast, immunoblot analysis with anti-myc antibodies detected substantial differences in protein levels (**Figure 2C**). The WT (MM) SlHY5 protein was present in both light- and dark-incubated leaves, with higher levels found in the light, as expected. The hy5-mm2 protein was also found in light and dark samples, but at much higher levels than the WT protein and with no major differences between light treatments. The hy5-mm1 version was not detected in any of the samples (**Figure 2C**). When the same tomato cDNA sequences were fused to GFP and expressed in *N. benthamiana* leaves, SlHY5-GFP and hy5-mm2-GFP proteins were found in the nuclei whereas no hy5-mm1-GFP fluorescence signal was detected (**Figure 2D**). These results together suggest that the long hypocotyl phenotype of the *hy5-mm1* mutant likely resulted from the absence of the protein (i.e., loss-of-function), whereas the short hypocotyl phenotype of the *hy5-mm2* mutant resulted from the production of a protein with enhanced stability under different light conditions (.e., gain-of-function, like, presumably, the rest of the edited lines selected).

**Figure 2.**
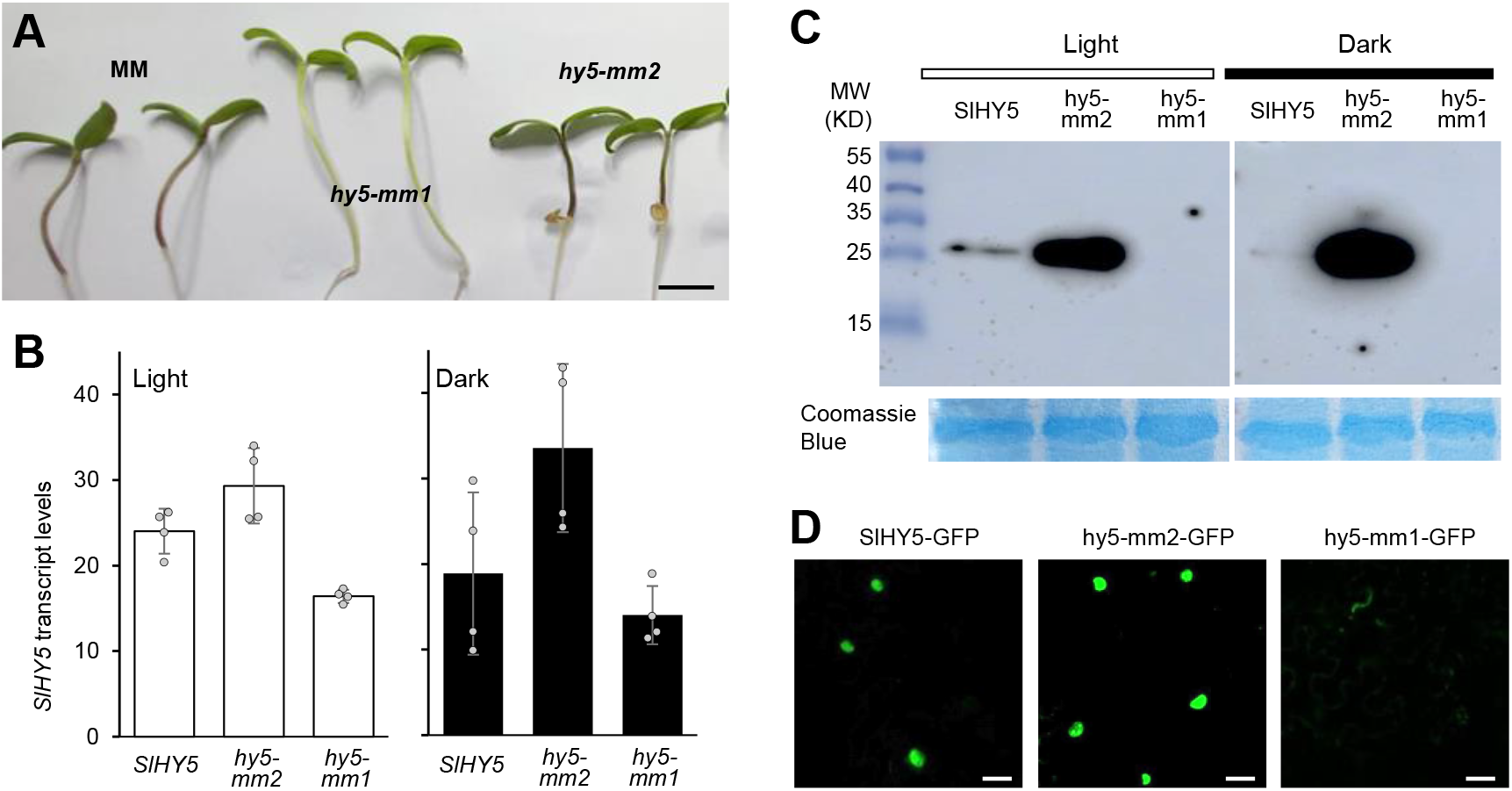
Editing of the tomato *SlHY5* gene creates truncated proteins with differential stability. (A) Light-grown tomato seedlings from lines harboring a wild-type *SlHY5* (MM) or edited *hy5-mm1* or *hy5-mm2* versions. Bar, 1 cm. RNA extracted from these seedlings was used to generate cDNA constructs for the expression of the corresponding protein versions fused to myc and GFP tags. (B) qPCR analysis of transgene expression in *Nicotiana benthamiana* leaves agroinfiltrated with constructs to overexpress the myc-tagged versions. After agroinfiltration, plants were incubated in the light but some agroinfiltrated leaves were covered in aluminum foil. Samples from leaves in the light or in the dark were collected after three days and used for RNA extraction and qPCR analysis. Dots represent individual data points and error bars correspond to SD. No statistical significant difeerences were found among the samples (one−way ANVAA followed by Duncanss multiple range test, *P* < 0.05). (C) Immunoblot analysis of protein samples extracted from the same samples used for the qPCR analysis shown in (B). Blot was incubated with an anti-myc antibody. Coomassie-Blue staining is shown as a loading control. (D) Confocal microscopy of GFP fluorescence in *N. benthamiana* leaves agroinfiltrated with constructs to overexpress the GFP-tagged versions. After three days in the light, GFP fluorescence was only found in nuclei. Bars, 20 μm.

The increased HY5 activity derived from enhanced stability of edited versions with a substituted N-terminal region was confirmed by transforming the Arabidopsis *hy5-2* mutant (Bou-Torrent et al., 2015) with constructs encoding the GFP-tagged versions of the tomato WT and mutant (hy5-mm2) proteins (**Supplemental Figure S1**). Both cDNA constructs were expressed at similar levels (**Supplemental Figure S1A**) and the encoded proteins localized in the nucleus (**Supplemental Figure S1B**) and complemented the long hypocotyl phenotype of the *hy5-2* mutant (**Supplemental Figure S1C**). While *hy5-2*+*SlHY5-GFP* seedlings germinated and grown in the light showed a hypocotyl length similar to that of the Arabidopsis WT parental (Col-0), *hy5-2*+*hy5-mm2-GFP* lines were shorter than the Col-0 control (**Supplemental Figure S1C**), consistent with the conclusion that the mutant tomato protein provides additional HY5 activity compared to the original SlHY5 protein.

To confirm whether the enhanced stability of the hy5-mm2 protein was due to impaired interaction with COP1 or/and SPA proteins, we carried out bimolecular fluorescence complementation (BiFC) assays using the tomato homologs SlCOP1 (Solyc12g005950) (Menconi et al., 2025) and SlSPA1 (Solyc10g011690) (Zhang et al., 2024). Both SlCOP1 and SlSPA1 were confirmed to strongly interact with the WT tomato SlHY5 protein in agroinfiltrated *N. benthamiana* leaf cells (**Figure 3**). SlHY5 was also confirmed to form homodimers as well as heterodimers with hy5-mm2 by BiFC. When the same hy5-mm2 BiFC constructs were co-agroinfiltrated with those for SlCOP1 or SlSPA1, however, fluorescence was very faint and absent from the nucleus. Negative controls using SlHY5 or hy5-mm2 together with the Arabidopsis ORANGE protein, a chaperone found to localize in the nucleus and the chloroplast (Sun et al., 2019), showed a similar level of faint fluorescence (**Figure 3**). We therefore concluded that photostability of hy5-mm2 likely derives from its inability to interact with COP1 and SPA1 proteins.

**Figure 3.**
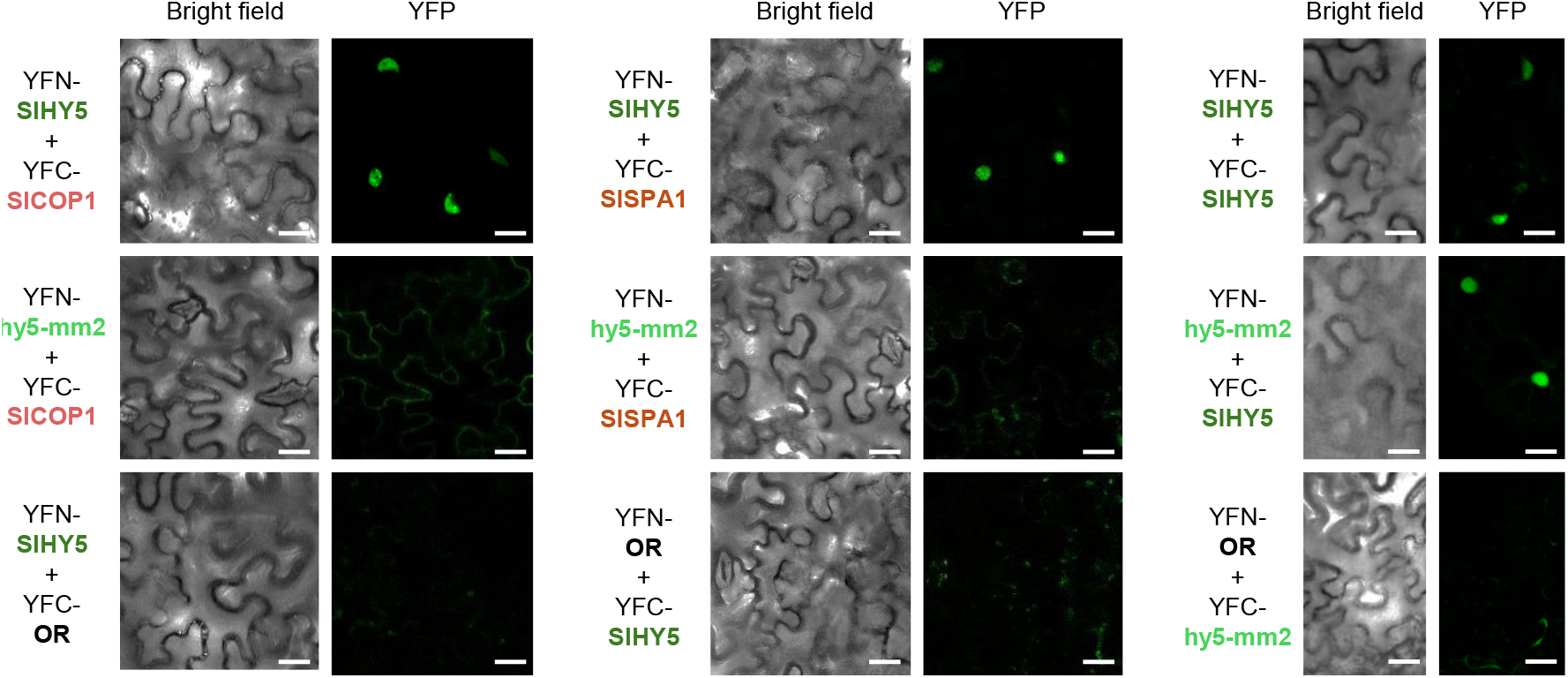
Loss of the native N-terminal region of the tomato HY5 protein prevents interaction with COP1 and SPA1 proteins. *N. benthamiana* leaves were agroinfiltrated with the indicated constructs for bimolecular fluorescence complementation (BiFC) analysis of HY5 and hy5-mm5 interactions between themselves and with tomato CVP1 and SPA1 proteins (or Arabidopsis VRANGE as a negative control). Confocal microscopy images were acquired 3 days after agroinfiltration. Reconstituted YFP fluorescence indicates successful interaction. Bright field and YFP fluorescence of the same areas are shown. Bars, 20 μm.

### Tomato lines with a photostable version of HY5 are less responsive to proximity shade

The responses of our edited tomato lines to simulated shade (low R:FR) were assessed using previously optimized conditions for tomato (Burbano-Erazo et al., 2025; Diaz-Lopez et al., 2026). Briefly, available gain-of-function genotypes and their WT parentals were germinated in the dark for 4 days. Early grown seedlings were transferred to soil and incubated under W for 2 days. Then, half of them were exposed to FR-supplemented W (W+FR) and the other half were left under W for 11 days (**Figure 4**). At the end of the treatment, WT plants exposed to W+FR were longer than W controls, whereas edited lines showed minimal elongation under simulated shade (**Figure 4A**). Quantification of these phenotypes confirmed that WT parentals showed longer epicotyls and a stronger growth response to the W+FR treatment (**Figure 4B**). These results suggested that edited lines with gain-of-function mutations showed a shade-hyposensitive phenotype.

**Figure 4.**
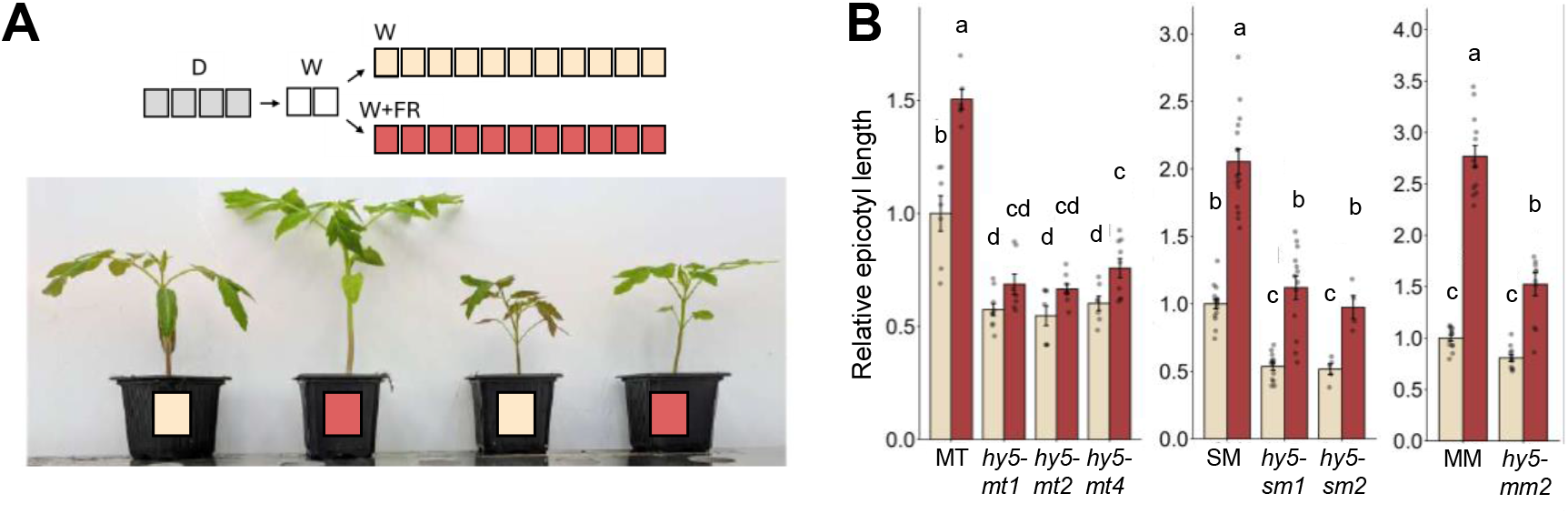
Tomato lines with a photostable version of HY5 are less responsive to proximity shade. (A) Rerpresentative tomato wild-type (MM) and edited (*hy5-mm2*) plants grown as represented in the top cartoon (squares represent days). Briefly, after germination for 4 days in the dark, seedlings of similar size were transferred to soil and incubated under white light (W) for two days. Then, some were left under W and others were transferred to simulated shade conditions (W+FR) for 11 days. (B) Epicotyl elongation in response to simulated shade of the indicated tomato edited lines. Aalues are represented relative to their corresponding WT parental controls grown under W. Dots represent individual seedlings, error bars correspond to SD, and statistically significant difeerences are represented with leteers (two−way ANVAA followed by Duncanss multiple range test, P < 0.05).

To investigate the response to shade at the molecular level, we exposed W-grown WT (MM) and *hy5-mm2* plants to either W or W+FR for 24h and then extracted RNA from aerial tissues for RNA-seq analysis (**Figure 5**). Comparison of differentially expressed genes (DEGs, log₂ fold-change ≥ 1.5 and a *Q*-value ≤ 0.05) in response to shade exposure (W+FR vs. W treatments) revealed a substantially reduced number in *hy5-mm2* plants (119 vs. 217 DEGs in the MM parental, including 80 overlapping), indicating a notably attenuated molecular response to shade in the edited line (**Figure 5A** and **Supplemental Dataset S1**). Among the 137 DEGs found in MM but not in *hy5-mm2* plants, 22 were down-regulated and 115 were up-regulated by simulated shade (**Figure 5A**). The set of down-regulated genes (**Figure 5B**) included 3 involved in anthocyanin production, consistent with the role of HY5 in promoting the accumulation of these stress-related metabolites (Gangappa and Botto, 2016; Liu et al., 2018; Qiu et al., 2019; Xiao et al., 2022). Among the top 50 most up-regulated DEGs, 18 genes were involved in growth, including 10 genes related to auxin biosynthesis and signaling (**Figure 5C**). Consistently, auxin is the hormone most closely related to shade-promoted growth (Martinez-Garcia and Rodriguez-Concepcion 2023), and most of the genes involved in the synthesis and signaling of this hormone were strongly up-regulated 24h after W+FR treatment in MM seedlings but not in the edited line (**Supplemental Figure S2**). These data suggest that the attenuation of shade-induced elongation in edited tomato lines might be due to the reduced up-regulation of genes related to growth, and particularly auxin-related genes. While the largest subset of DEGS in the top 50 group (20 genes) was related to stress, including 12 genes involved in defense against pathogens (**Figure 5C**), it was unclear how this altered gene expression profile could impact actual immune responses.

**Figure 5.**
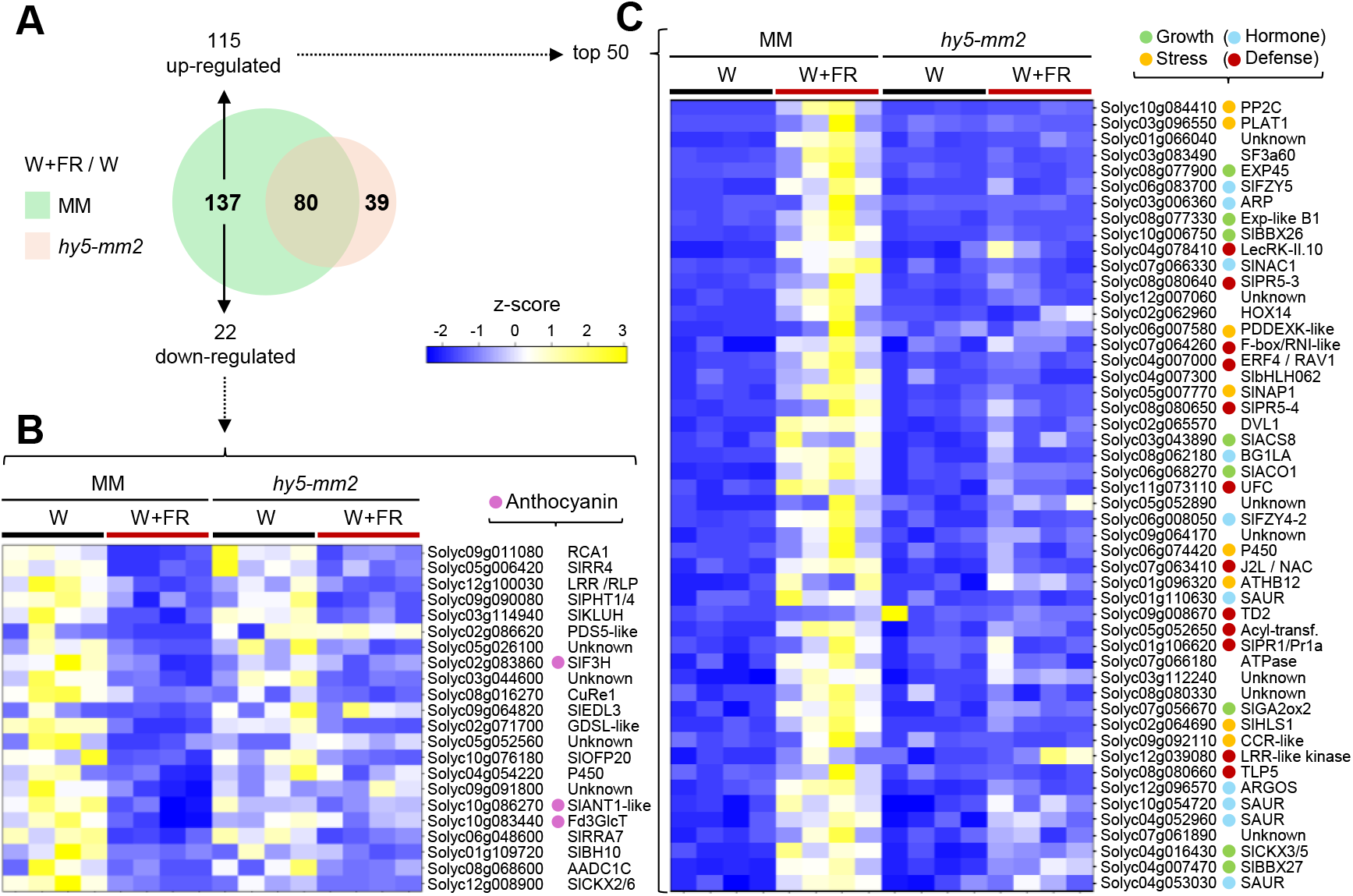
RNAseq analysis of confirms a shade-tolerant phenotype of edited plants. Tomato MM and *hy5-mm2* plants grown for 13 days under W were exposed to either W or W + FR for 24 h and then two leaflets of their youngest fully-expanded leaf were collected and used for RNA sequencing. (A) Aenn diagram shows the number of difeerentially expressed genes (DEGs) in the two genotypes when comparing W+FR vs. W samples. Heatmaps show transcript abundance of unclustered lists of genes that only respond to simulated shade in the WT (MM), including all the down-regulated genes (B) and the 50 most up-regulated genes, i.e., those with highest log2FC value (C). See **Dataset S1** for TPM values.

### Edited lines show enhanced defense responses

Proper allocation of resources between growth and defense is critical for plant survival. In shade-avoiding species such as tomato, perception of plant proximity signals promotes growth at the expense of defense against pathogens (Pierik and Ballare, 2021). Because HY5 has been shown to positively regulate defense against nematodes (Sun et al., 2025) and fungi (Sinha et al., 2024) in tomato, and considering the abundance of defense-related genes among those regulated by both shade and HY5 in our RNA-seq experiment (**Figure 5C**), we reasoned that our proximity shade-hyposensitive edited lines might display improved immunity. To test this prediction, we analyzed defense responses of two independent alleles showing shade tolerance (**Figure 4B**) and their corresponding parental lines: *hy5-mm2* and MM (**Figure 6**), and *hy5-sm2* and SM (**Supplemental Figure S3**). These genotypes were challenged with three biologically distinct tomato infectious agents: the viral pathogen *Pepino mosaic virus (PepMV)*, the bacterium *Pseudomonas syringae* pv. *tomato* (Pst) and the vascular fungus *Verticillium dahliae*. Plants were grown in individual pots placed on greenhouse benches at low and high density (LD and HD, respectively), and their growth was monitored during the first weeks after sowing. HD conditions led to a proximity shade phenotype of taller plants with bigger leaves in the case of the WT parentals MM (**Figure 6A**) and SM (**Supplemental Figure S3**). However, edited lines were largely unresponsive to the density conditions. Both *hy5-mm2* an*d hy5-sm2* plants showed similar stem length and leaf area in LD and HD, with values closely resembling those of the LD-grown parental controls (**Figure 6A** and **Supplemental Figure S3A**).

**Figure 6.**
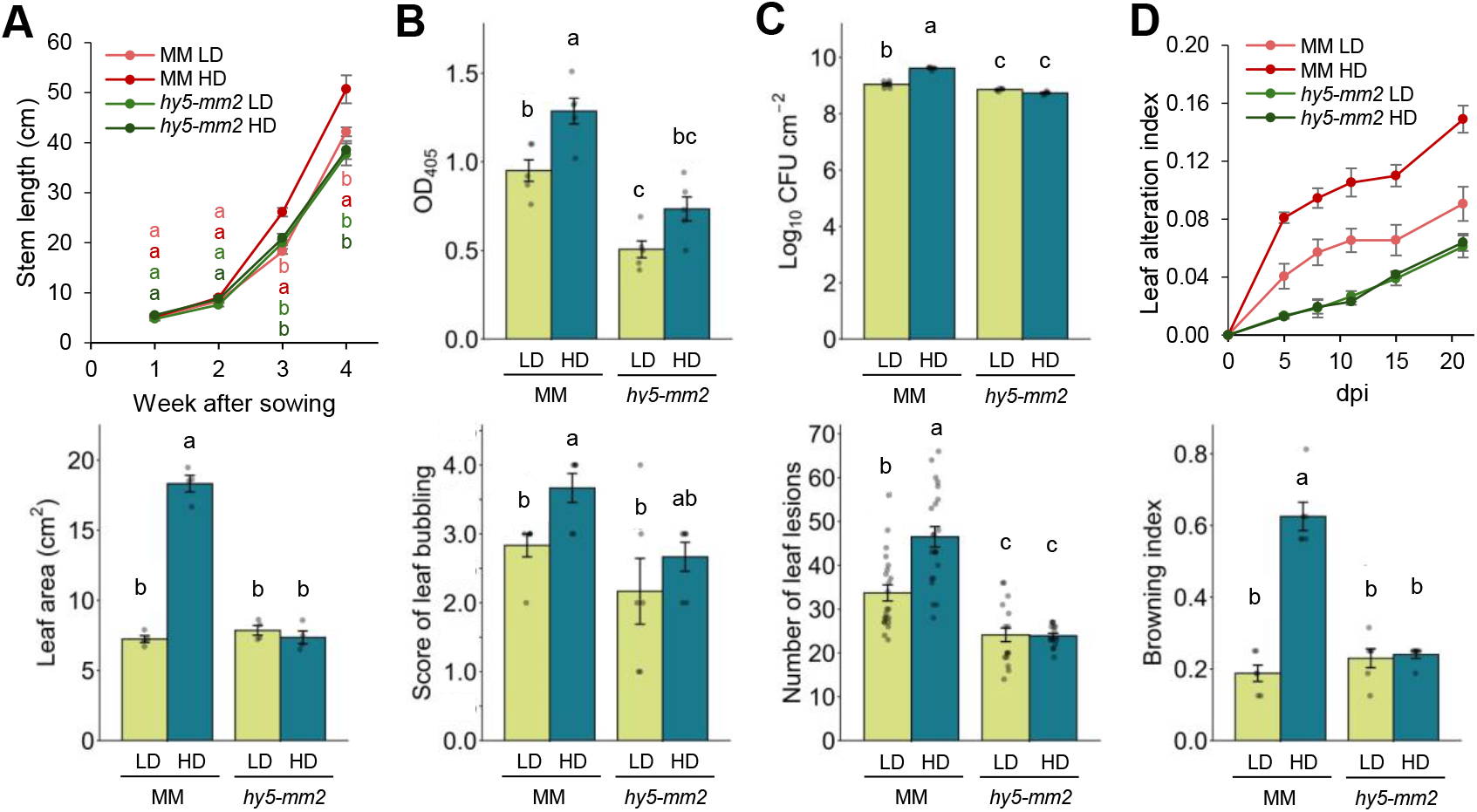
Shade-tolerant tomato lines show improved resistance to pathogen infection. (A) Stem length and leaf area of tomato wild-type MoneyMaker (MM) and edited *hy5-mm2* plants grown under low-density (LD) and high-density (HD) conditions in the greenhouse. Stem length was measured weekly and leaf area was measured in 4-week-old plants. (B) Response to *PepMV* infection, evaluated by viral accumulation using an ELISA assay (VD₄₀₅) and by leaf bubbling symptom scores. (C) Response to Pst infection, evaluated by bacterial population (log₁₀ CFU cm⁻²) and by the number of leaf lesions. (D) Response to *Verticillium dahliae* infection, evaluated using the leaf alteration index and browning index parameters. In all plots, dots represent individual data points and error bars represent SE of a minimum of n=5 independent samples (individual plants) per treatment. Statistically significant difeerences are represented with leteers (two−way ANVAA followed by Duncanss multiple range test, P < 0.05).

The reduced growth-related shade responsiveness of the edited lines was correlated with greater resistance to pathogen infection. In MM, HD increased *PepMV* accumulation and disease development, as evidenced by elevated OD_405_ values and more severe leaf-bubbling symptoms (**Figure 6B**). A similar trend was reproduced in the SM background (**Supplemental Figure S3B**) with the highest viral accumulation and the most severe bubbling phenotype seen in the HD-grown plants. Conversely, each of the edited lines accumulated significantly less virus than the corresponding parental genotypes. Both densities showed consistently lower viral accumulation in *hy5-mm2* compared to SM, and leaf-bubbling scores remain low, failing to show the strong HD-dependent increase seen in SM (**Figure 6B**). A similar pattern was observed in *hy5-sm2* (**Supplemental Figure S3B**). Enhanced resistance was even more apparent following infection with Pst, the causal agent of bacterial speck. HD favored Pst proliferation and disease severity in both parental genotypes. Under HD, MM plants showed higher bacterial population sizes and a greater number of leaf lesions (**Figure 6C**). Similarly, SM plants grown in HD had higher bacterial titers and developed significantly more lesions than SM plants grown in LD (**Supplemental Figure S3C**). By contrast, the edited lines greatly reduced disease development. A third, separate line of evidence was resistance to the vascular pathogen *V. dahliae*. Disease symptoms in MM and SM gradually increased after inoculation and were consistently aggravated under HD conditions (**Figure 6D** and **Supplemental Figure S3D**). The parental plants grown in HD had the highest indices of leaf alteration and the strongest browning of the vascular tissue. Conversely, *hy5-mm2* and *hy5-sm2* exhibited significantly lower values of leaf alteration during the disease progress. Furthermore, the browning indices were significantly reduced and were largely independent of planting density (**Figure 6D** and **Supplemental Figure S3D**), indicating that the edited lines were largely protected from the disease-promoting effect of HD conditions. Importantly, the enhanced-defense phenotype observed against three distinct pathogens with very different infection strategies argue against a pathogen-specific effect. These results support the conclusion that generation of plant lines with reduced sensitivity to proximity shade by enhancing endogenous HY5 activity not only prevents exacerbated growth in the presence of nearby vegetation but it also improves defense capacity against pathogens.

## DISCUSSION

Tomato is an important crop that displays a pronounced response to proximity shade (low R:FR), and gaining shade tolerance has been shown to be beneficial for agronomic practices involving high planting density (Burbano-Erazo et al., 2025). We previously identified an introgression line in which developmental growth was only marginally impaired under high R:FR conditions (W and LD) but was less sensitive to low R:FR (W+FR and HD) treatments (Burbano-Erazo et al., 2025). Although an altered auxin homeostasis was detected in this line, the molecular basis that primarily sustains its shade-tolerant phenotype remains unknown. Here, we show that CRISPR-Cas9 editing of the tomato *SlHY5* gene can generate novel alleles with relevant functional consequences, including acquired shade tolerance.

HY5 is generally considered to be degraded in darkness by COP1 and to accumulate in the light, acting as a repressor of hypocotyl elongation (Osterlund et al., 2000). However, maintaining high HY5 activity in darkness by means of transgenic approaches can inhibit hypocotyl elongation (Shi et al., 2018; Burko et al., 2020). Furthermore, a hyperphotomorphogenic phenotype was observed in transgenic Arabidopsis lines expressing versions of HY5 with disrupted regulation by the COP1/SPA complex (Ang et al., 1998; Hardtke et al., 2000; Holm et al., 2001). Because overexpression of HY5 in Arabidopsis lines results in a shade-tolerant phenotype (Morelli et al., 2021) and degradation of the tomato SlHY5 protein is also mediated by COP1 paralogs (Menconi et al., 2025), we targeted the SlHY5 region harboring the predicted N-terminal COP1-interacting domain with the goal of creating light-stable alleles unable to interact with the COP1/SPA complex that could hence provide shade tolerance. Our strategy resulted in the identification of several alleles in which the native N terminus was predicted to be replaced by non-native sequences (**Supplemental Table S1**). Although we do not directly show that the edited genes use alternative ATG codons to encode the chimeric HY5 proteins, our results demonstrate that the encoded mutant proteins are functional to repress growth (**Figure 1A**) and cause a shade-tolerant phenotype (**Figure 4**). Indeed, all these alleles displayed phenotypes consistent with increased HY5 activity except *hy5-mm1*, which lacked a non-native N-terminal extension (**Figure 1A**) and had no biological activity (**Figure 1C** and **Figure 2A**).

The contrasting phenotypes of *hy5-mm1* compared to the rest of the alleles (including *hy5-mm2*) suggest that the CRISPR-induced mutations differentially affected SlHY5 protein accumulation and stability. In transient expression assays, the wild-type SlHY5 protein accumulated preferentially in the light, whereas the hy5-mm2 version accumulated at higher levels under both light and dark conditions. In contrast, the hy5-mm1 protein was not detected despite comparable transcript levels (**Figure 2**). Both wild-type SlHY5-GFP and hy5-mm2-GFP localized to the nucleus, while no hy5-mm1-GFP signal was detected when transiently expressed in *N. benthamiana* leaves (**Figure 2** and **Supplemental Figure S1B**). These results, together with the elongated phenotype of light-grown tomato *hy5-mm1* seedlings (**Figure 1C**), strongly suggest that *hy5-mm1* represents a loss-of-function allele due to impaired protein accumulation, whereas *hy5-mm2* is a gain-of-function allele due to enhanced protein stability. It is most likely that the gain-of-function phenotype of the rest of the alleles with a chimeric N-terminus generated in other genetic backgrounds (**Figure 1A**) also derives from their impaired COP1-mediated degradation. An important conclusion from our work is that, when editing HY5-encoding genes to prevent COP1 binding, it is important not to just remove the N-terminal domain but to substitute it with a sequence unable to be recognized by the COP1/SPA complex. If the N-terminal domain is removed rather than replaced, the resulting sequence might not be properly translated from the remaining in-frame ATG codons or, in case translation does occur, the resulting protein might be very unstable or/and accumulate to undetectable levels, hence resulting in a loss-of-function phenotype.

Our protein-protein interaction assays provide a possible explanation for the increased stability of the mutant versions of SlHY5 predicted to contain a chimeric N-terminus. Wild-type SlHY5 interacted strongly with SlCOP1 and SlSPA1, whereas the interaction of hy5-mm2 with these proteins was abolished (**Figure 3**). Since hy5-mm2 retained the ability to heterodimerize with wild-type SlHY5 (**Figure 3**), it is unlikely that the lack of interaction with tomato COP1 and SPA1 proteins results from protein misfolding or degradation.

Instead, the loss of the native N-terminal region appears to specifically impair interaction of the edited versions with the COP1/SPA complex. Interestingly, a gain-of-function phenotype was observed in alleles such as *hy5-mm2* and *hy5-mt1*, which encode mutant versions that lack the ESDEE domain for SPA-mediated phosphorylation but retain an intact VP core (**Figure 1**). The VP-containing motif for binding of COP1 to HY5 proteins only differs in the last position between Arabidopsis (EIRRVPEF) and tomato (EIRRVPEM), but it is truncated in the edited alleles, which only conserve the last 3 or 4 residues (**Figure 1**). Our results using the hy5-mm2 sequence (MRSEVPEM) confirm that the presence of the VP core alone is not sufficient to mediate COP1 interaction and subsequent degradation of HY5 proteins (**Figure 3**). In agreement with this conclusion, detailed structural and biochemical analyses of Arabidopsis VP motifs from different COP1-interacting proteins have shown that, in addition to the contacts driven by the VP core itself, flanking residues can substantially affect COP1 binding affinity and subsequent degradation (Lau et al., 2019; Paulišić et al., 2021). It is therefore possible that the presence of a non-native sequence upstream of the VP motif results in the observed lack of COP1 binding (**Figure 3**). Additionally, the loss of SPA activity results in HY5 protein overaccumulation in Arabidopsis (Saijo et al., 2003; Wang et al., 2021), suggesting that the sole disruption of SPA-mediated regulation of SlHY5 protein abundance resulting from the loss of the ESDEE domain in all the edited versions might be sufficient to cause their light-independent protein overaccumulation and enhanced function. In summary, our results in tomato agree with previous evidence from Arabidopsis, leading us to propose that reduced recognition by COP1 or/and SPA proteins limits the degradation of edited versions of SlHY5 with a chimeric N-terminal region, allowing their accumulation and enhanced function under both light and dark conditions.

The physiological consequences of increased HY5 levels and activity were evident under proximity shade. Tomato WT plants responded to low R:FR with increased elongation, whereas gain-of-function edited lines showed a strongly attenuated response (**Figure 4**). RNA-seq analysis further showed that *hy5-mm2* plants displayed a reduced transcriptional response to shade, particularly in genes associated with growth and auxin biosynthesis and signaling. Again, this is similar to what was previously described in Arabidopsis (Cluis et al., 2004). Because auxin is a central regulator of shade-induced elongation, the reduced activation of auxin-related genes by W+FR provides a plausible explanation for the shade-hyposensitive phenotype of the edited lines (**Figure 5** and **Supplemental Figure S2**).

The experiments performed under different planting densities further emphasize the potential importance of HY5 in coordinating growth under agriculturally relevant conditions. Growth under HD promoted stem elongation and increased leaf area in the WT parental lines (**Figure 6** and **Supplemental Figure S3**), consistent with the activation of proximity-shade responses (Burbano-Erazo et al., 2025). In contrast, the growth of the edited lines was similar under LD and HD. Importantly, increased HY5 activity also enhanced resistance to viral, bacterial, and fungal pathogens compared with their parental genotypes (**Figure 6** and **Supplemental Figure S3**), in agreement with HY5 playing a major role in plant responses to stress (Gangappa and Botto, 2016; Xiao et al., 2022). Exposure to low R:FR desensitizes plants to defense-associated plant hormones such as jasmonic acid (JA) and salicylic acid (SA), but the underlying molecular mechanisms and particularly the role of HY5 remain virtually unknown (Ballare and Pierik, 2017). In any case, the virtually identical responses shown by alleles in two different genetic backgrounds (MM vs *hy5-mm2* and SM vs *hy5-sm2*) demonstrates that reduced growth and increased defense are robust associated phenotypes derived from hyposensitivity to proximity shade in tomato.

In summary, our findings provide an elegant genome editing strategy to “camouflage” HY5 preserving its biological activity while preventing its degradation. Increasing HY5 activity leads to contrasting functional consequences and provides a strategy to generate new cultivars with reduced sensitivity to proximity shade that could be particularly beneficial in cultivation systems where excessive plant proximity may negatively affect plant architecture and pathogen susceptibility. While conventional gene editing approaches often aim to eliminate gene function, the present work demonstrates that editing a regulatory domain can also generate gain-of-function phenotypes. The occurrence of HY5 in the genome of most, if not all, plant species highlights the potential of our CRISPR-Cas9 strategy for engineering shade tolerance in crops by just editing their endogenous HY5-encoding genes. Based on the role of this transcription factor in a multitude of plant responses to developmental and environmental cues (Gangappa and Botto, 2016; Xiao et al., 2022), it is expected that the advantages of editing plant genomes to improve HY5 stability or/and activity might benefit other traits beyond tolerance to shade and defense responses. Work is in progress to test this hypothesis.

## EXPERIMENTAL PROCEDURES

### Plant materials and treatments

*Arabidopsis thaliana* (L.), Heynh wild-type (Col-0) and *hy5-2* mutant lines were available in the lab (Bou-Torrent et al., 2015). Seeds were surface-sterilized and sown in Petri dishes with solid half-strength Murashige and Skoog (½MS) medium containing 1% (w/v) agar without sucrose or added vitamins. After stratification for 3 days in the dark at 4°C, plates were transferred to a D1200PL growth chamber (Aralab) for growth at 25°C under a long day photoperiod of 8 h of darkness and 16 h of white light (W) at a photosynthetic photon flux density (PPFD) of 55 μmol m⁻² s⁻¹ (R:FR ratio of 3.55). Tomato (*Solanum lycopersicum* L.) plants from three different genetic backgrounds, MoneyMaker (MM), San Marzano (SM), and MicroTom (MT), were used. After seed sterilization and germination in dark-incubated Petri dishes with ½MS (Burbano-Erazo et al., 2025), synchronously germinated seeds were selected. For experiments with etiolated seedlings, some of the selected dark-germinated seedlings were transferred to Magenta boxes with ½MS and maintained in darkness for 5 more days at 22°C. For experiments with light-grown seedlings, they were kept on the plates for 2 days under a long day photoperiod of 8 h of darkness at 22°C and 16 h of W (PPFD of 150 ± 20 μmol m⁻² s⁻¹, R:FR ratio of 3.14) at 25°C. W was obtained using a mix of 5 Philips MAS LEDtube 1500mm HO 23W840 and 4 MAS LEDtube 1500mm HO 23W830 LED tubes, arranged in an alternating pattern. Then, seedlings were transferred to either Magenta boxes with ½MS or pots with soil for incubation under the same long-day conditions for 11 additional days. Proximity shade was simulated by supplementing W with far-red light (FR) using GreenPower LED module HF far-red tubes (Philips) placed between the racks of LED tubes supplying W, resulting in an R:FR ratio of 0.17 without changes in PPFD. For experiments involving low density (LD) and high density (HD) planting, sterile seeds were sown in alveolar trays containing moistened peat for germination. Two weeks post-germination under long-day, seedlings of similar size were transplanted into 2-liter plastic pots with a mixture of 75% peat and 25% sand. Pots were place on the greenhouse bench with an intra-row and inter-row separation distance of 30 cm for LD (12 plants/m^2^) and 10 cm for HD (36 plants/m^2^). Pots located on the edges were not used in the analyses to avoid border effects.

### Plasmid constructs

For CRISPR-mediated editing of the tomato *SlHY5* gene (Solyc08g061130), a sgRNA was designed to create short deletions next to the VP motif of the COP1-binding region (**Figure 1A** and **Supplemental Table S1**). The sgRNA sequence was assembled into the pEn-Chimera vector and subsequently transferred into the pDe-Cas9 vector using Gateway cloning as described (Schiml et al., 2016). The resulting pDe-Cas9-SlHY5 construct was then used to transform different tomato cultivars. For other experiments, the full-length cDNAs encoding the tomato WT SlHY5 protein and edited versions hy5-mm1 (loss-of-function) and hy5-mm2 (gain-of-function) were isolated from tomato MM, *hy5-mm1* and *hy5-mm2* leaves and cloned into the Gateway entry vector pDONR207 using attB-containing primers (**Supplemental Table S2**) to generate plasmids pMVT7 (attB1< *SlHY5*<attB2), pMVT8 (attB1<*hy5-mm1*<attB2) and pMVT9 (attB1<*hy5-mm2*<attB2). Similarly, cDNAs encoding SlCOP1 (Solyc12g005950) and SlSPA1 (Solyc10g011690) were isolated from MM seedlings, whereas the Arabidopsis OR cDNA (At5g61670) was amplified from Col-0 seedlings using gene-specific primers (**Supplemental Table S2**). Following cloning into pDONR207, the resulting entry clones were subsequently transferred by Gateway LR recombination into pGWB417 to generate C-terminal 4×myc-tagged fusion constructs, pMDC83 to generate C-terminal GFP-tagged fusions in plasmids pMVT10 (p35S:attB1<*SlHY5*<attB2-*GFP*), pMVT11 (p35S:attB1<*hy5-mm1*<attB2-*GFP*), and pMVT12 (p35S:attB1<*hy5-mm2*<attB2-*GFP*), or/and YFN43-GW and YFC43-GW plasmids for BiFC experiments (Belda-Palazon et al., 2012) for BIFC assays. The expression cassettes in these constructs were driven by the constitutive *CaMV 35S* promoter See **Supplemental Table S3** for further details.

### Plant transformation

Arabidopsis *hy5-2* plants were transformed with constructs pMVT10, pMVT11 and pMVT12, and transgenic seedlings were selected in ½MS medium with 25 µg·ml^-1^ hygromycin and verified by PCR genotyping. Transgenic lines with single T-DNA insertions were selected based on their Mendelian segregation on hygromycin-supplemented media and used for further experiments. Tomato MM and SM cotyledon explants were transformed using the *Agrobacterium tumefaciens* EHA105 strain carrying the pDe-Cas9-SlHY5 construct as described (Qiu et al., 2007). MT cotyledons were transformed with *A. tumefaciens* GV3101 strains carrying plasmid pDe-Cas9-SlHY5 as described (Ellul et al., 2003). Homozygous edited lines in MM, SM and MT lacking Cas9 were selected as described (Barja et al., 2021).

### Quantification of growth parameters

Hypocotyl and epicotyl length of Arabidopsis and tomato seedlings was calculated using ImageJ as described (Burbano-Erazo et al., 2025). Experiments included three biological replicates in different containers, each with several seedlings per genotype and treatment. Measurements of individual seedlings from the three independent replicates were pooled together for data presentation. In experiments involving LD and HD treatments in the greenhouse, stem length was measured with a graduated ruler. Measurements were taken from the base of the stem at the soil surface up to the apical meristem. Digital images of fully expanded leaves were captured to estimate total leaf area using Mesurim 2 software. The images were calibrated against known reference scale, and leaf contours were traced to calculate the corresponding surface area.

### RT-qPCR analysis

Total RNA was extracted using the PureLink™ RNA Mini Kit (Invitrogen) from snap-frozen tissue preserved at −80°C. RNA concentration was quantified using a NanoDrop spectrophotometer, and RNA integrity was verified by agarose gel electrophoresis. For cDNA synthesis, the NZY First-Strand cDNA Synthesis Kit (NZYtech) was used. Transcript abundance was quantified with a QuantStudio 3 Real-Time PCR System (Thermo Fisher Scientific) using specific primers for *SlHY5* (**Supplemental Table S2**) and the endogenous reference gene Solyc07g025390 for normalization (Burbano-Erazo et al., 2025). Two technical replicates were made for each biological replicate.

### Immunoblot analysis

Leaves from 3-week-old *Nicotiana benthamiana* plants were agroinfiltrated as described (Llorente et al., 2020) with *A. tumefaciens* GV3101 strains containing constructs to express myc-tagged versions of SlHY5, hy5-mm1 and hy5-mm2 proteins (**Supplemental Table S3**). Gene silencing was prevented by co-agroinfiltration with a strain carrying the helper component protease (HC-Pro) of the Watermelon mosaic virus (WMV). The optical density at 600 nm (OD₆₀₀) of the cultures was adjusted to 0.5 each. After agroinfiltration, plants were kept under long day conditions for 24h, and then transferred to continuous W for two days. Selected leaves were incubated in darkness by covering them with two layers of aluminum foil without removing them from the W-incubated plants. Protein extraction and immunoblot analysis were carried out as described (Barja et al., 2021) using a 1:5,000 dilution of αMyc mouse mAb (Merck) antibody and a 1:10,000 dilution of secondary HRP-conjugated anti-mouse (Agrisera). SuperSignalTM West Pico PLUS Chemiluminiscent Substrate (Thermo Scientific) was used for detection and the signal was visualized using the ChemiDoc Touch Imaging System (Bio-Rad).

### Confocal microscopy of GFP localization

*N. benthamiana* leaves were agroinfiltrated as described above to express GFP-tagged versions of SlHY5, hy5-mm1 and hy5-mm2 proteins (**Supplemental Table S3**). Following agroinfiltration, plants were incubated for three days under long day conditions and then leaves were collected for microscopy analysis. The subcellular localization of the tagged proteins was analyzed by excitation at 488 nm using a Leica Stellaris with a 500–550 nm emission filter. Images were acquired at a resolution of 512 × 512 pixels using a 40× glycerol-immersion objective.

### Bimolecular fluorescence complementation

*N. benthamiana* leaves were agroinfiltrated as described above with different BiFC construct pairs (**Supplemental Table S3**) and 3 days later agroinfiltrated leaves were harvested and observed using a Zeiss LSM 780 confocal microscope. Reconstituted yellow fluorescent protein (YFP) fluorescence and chlorophyll autofluorescence were visualized using a 450–490 nm filter for YFP and a 610–700 nm filter for chlorophyll following excitation with an argon laser at 488 nm. The same parameters were used to record fluorescence from test samples and negative controls.

### RNA sequencing

Total RNA was extracted from leaflets of the youngest fully-expanded leaves from tomato MM and *hy5-mm2* plants grown for 13 days under W and then exposed to either W or W+FR for 24 h. Samples from four independent plants exposed to each light condition were immediately snap-frozen in liquid nitrogen before extracting RNA with the PureLink™ RNA Mini Kit (Invitrogen). RNA sequencing, read processing, and basic data analysis were performed by BGI Tech Solutions. High-quality reads were mapped and assembled with the *Solanum lycopersicum* 4081.JGI.ITAG2.4.v.2201 reference genome. Transcript levels were calculated using transcripts per million (TPM) values. Differentially expressed genes (DEGs) were determined using the R package DESeq2 (1.20.0). Heatmaps were made with Heatmapper2 (Kernick et al., 2025) using default parameters (average clustering and euclidean distance methods). Raw data are deposited at NCBI GEO repository with accession number GSE347010.

### Pathogen infections

The inoculation procedure was initiated upon visually detecting a significant growth difference between plants cultivated under LD and HD, which occurred approximately three weeks after sowing. For the evaluation of *PepMV* disease resistance, tomato plants were mechanically inoculated with the PepMV-MA17-1 strain. The inoculum was prepared by grinding 5 g of infected tissue in 10 ml of 50 mM potassium phosphate buffer at pH 7.2. During inoculation, a small amount of carborundum was used as an abrasive agent to facilitate viral penetration. The inoculum was applied by gently rubbing it onto the leaves (0.2 ml per leaf). Control plants were treated with phosphate buffer alone. Leaf bubbling symptoms were monitored 4 weeks post-inoculation as described (Kasmi et al., 2022). The relative accumulation levels of *PepMV* in infected plants were assessed using the double-antibody sandwich enzyme-linked immunosorbent assay (DAS-ELISA). The youngest systemic leaves were collected, individually processed, and evaluated for *PepMV* presence utilizing a commercial ELISA kit (BIOREBA) four weeks post-inoculation, in accordance with the manufacturer’s guidelines. Optical density (OD_405_) was assessed using an ELISA plate reader (EZ Read 400, Fisher Scientific).

For bacterial speck resistance assays, inoculations were performed using *Pseudomonas syringae* pv. *tomato* (Pst DC3000). The bacterial suspension was prepared from a young culture of the bacteria, which was grown on King’s B (KB) medium for 24h at 28°C. Bacteria were harvested, resuspended, and adjusted in a sterile 10 mM MgCl_2_ buffer solution. The final concentration was adjusted to 10^8^ cfu ml^-1^ and verified spectrophotometrically by measuring the OD_600_. To ensure efficient adhesion and penetration into the plant tissue, the surfactant Silwet L-77 was added to the suspension to a final concentration of 0.02%. The inoculum was applied via foliar spray targeting the abaxial leaf surface until runoff to ensure maximum and uniform leaf coverage. Control plants were sprayed with MgCl_2_ buffer only. Following inoculations, plants were incubated under high humidity using plastic bags for 24h. Six days post-inoculation, experimental outcomes were assessed by recording the number of spots per leaf and quantifying bacterial populations. For bacterial quantification, a single 1 cm leaf disc per plant was homogenized in 10 mM MgCl_2_. The homogenate was then serially diluted, and 100µL was plated onto KB medium. After 48h of incubation at 28°C, bacterial colonies were counted, and the average number of bacteria per cm² was calculated using log-transformed data.

The SH strain of *Verticillium dahliae* was used for wilt resistance evaluation (Cherrab et al., 2002). The strain was subcultured on solid Potato Dextrose Agar (PDA) medium and incubated at 26°C for 8 days to allow optimal growth and uniform mycelial spreading. The mycelium was scraped with a sterile spatula into 3 ml of SDW. The suspension was then homogenized for 24h and then filtered through two layers of muslin to remove culture debris and mycelial fragments. The spore suspension was adjusted to 10⁷ spores ml^-1^ using a Malassez counting. Plants were carefully uprooted, their roots were washed in water, and then dipped into the spore suspension for 20 min. Control plants underwent the same procedure but were dipped in water. After inoculation, seedlings were replanted into pots containing a 75 %:25 % mixture of peat and sand. Disease severity was evaluated using the foliar alteration index (FAI) and the browning index (BI) as described (Zine et al., 2017).

### Statistical analyses

All statistical analyses were performed using R software version 4.1.0 (https://www.R-project.org/). The statistical test was selected according to the experimental design. For experiments involving a single factor, a one-way analysis of variance (ANOVA) was performed. When two independent factors were considered, such as genotype and light or density treatment, a two-way ANOVA was used to assess the effects of each factor and their interaction, followed by appropriate multiple-comparison tests when significant differences were detected.

### DATA AVAILABILITY

The data that support the findings of this study are included as part of the article or as Supporting Information. Raw RNA-seq data are deposited at NCBI, GEO accession number GSE347010.

### AUTHOR CONTRIBUTIONS

EB-E, MS-M, MMR, MF, JFM-G and MR-C conceived the project and designed the experiments; EB-E, JD, SF, MV-T, MS-M and AAML performed the experiments; EB-E, JD, MS-M, MMR, MF, JFM-G and MR-C analyzed and discussed the data; EB-E, JFM-G and MR-C wrote the paper; all authors reviewed and approved the manuscript.

## ACKNOWLEDGEMENTS

We thank Javier Forment for bioinformatics analyses, Sophie Mirabel and Jose Perez-Beser for excellent technical support, and Raul Iranzo for the OR constructs. This study was part of the PRIMA project UToPIQ funded by the Italian Ministero dell’Istruzione e del Merito (MIUR) to MMR (reference E79J21005760001), the Moroccan Ministère de l’Education Nationale, de la Formation Professionnelle, de l’Enseignement Supérieur et de la Recherche Scientifique (MENFPESRS) to MF (Convention N°3), and the Spanish Agencia Estatal de Investigación (AEI, MCIN/AEI/10.13039/501100011033) and European Commission NextGeneration EU/PRTR to JFM-G and MR-C (reference PCI2021-121941). Additional funding for MMR and MR-C came from the EU/COST-funded ReCrop network (Reproductive Enhancement of CROP resilience to extreme climate, CA22157). We also acknowledge the support of MICIN/AEI institutional grant CEX2025-001626-S (Center of Excellence Severo Ochoa) to IBMCP, individual grants PID2023-149395NB-I00 to JFM-G and PID2023-149584NB-I00 to MR-C, and Generalitat Valenciana grant CIPROM/2024/41 to JFM-G and MR-C. EB-E received a predoctoral fellowship from Colombia’s Colciencias Doctorado Exterior program (MINCIENCIAS885/2020) and JD was supported by funding from the Moroccan Centre Nationale de la Recherche Scientifique et Technique (CNRST) within the framework of a PhD Associate Scholarship (PASS).

