## Supplemental data for "A photostable version of HY5 confers tolerance to proximity shade and improved defense responses in tomato"

### SUPPLEMENTAL FIGURES

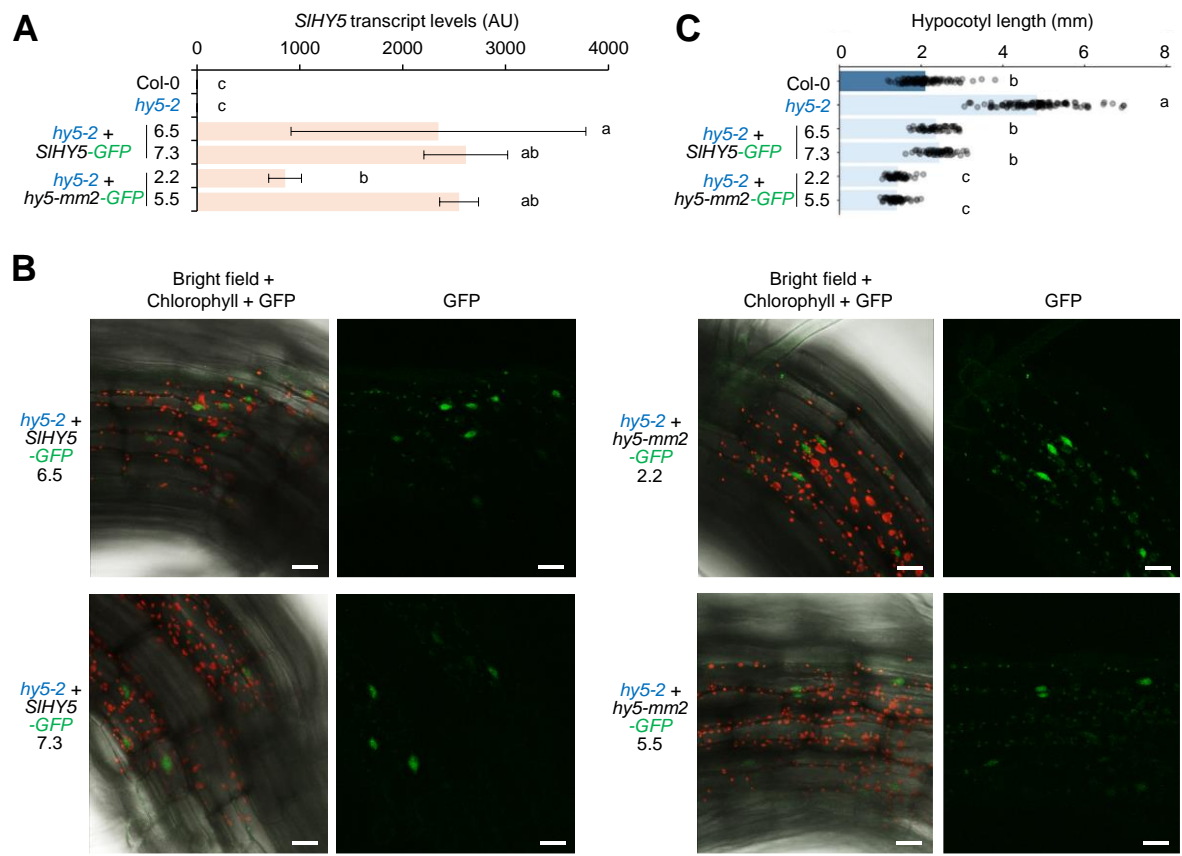

**Figure S1. Complementation of the Arabidopsis *hy5-2* mutant with tomato HY5 versions.** Mutant *hy5-2* seedlings were transformed with constructs to overexpress the WT *SlHY5* or the edited *hy5-mm2* versions fused to GFP. After germination, they were grown for a week under long-day conditions. (A) qPCR analysis of *SIHY5-GFP* and *hy5-mm2-GFP* transgene expression. Error bars correspond to SD of n=3 independent replicates. (B) Confocal microscopy of GFP fluorescence in the hypocotyl of the indicated lines. Left panels show bright field, chlorophyll autofluorescence (in red) and GFP fluorescence (in green), whereas right panels show only GFP signal in the same field. Bars, 20 μm. (C) Hypocotyl length in the indicated lines. In both plots, letters represent statistically significant differences (one-way ANOVA followed by Duncan's multiple range test, $P < 0.05$).

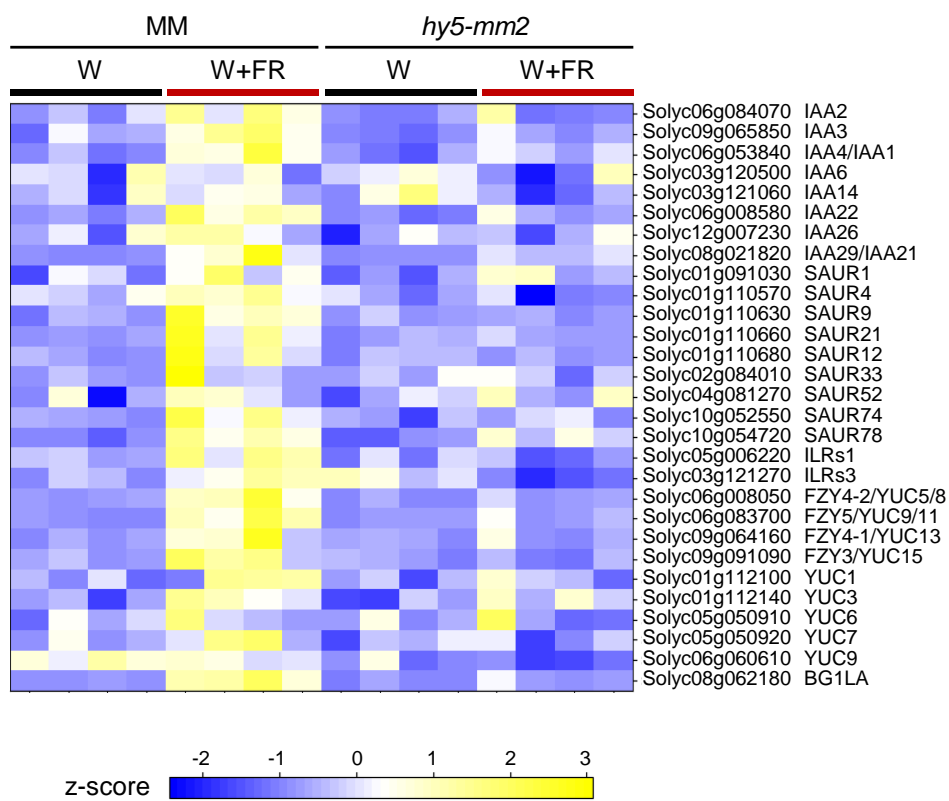

**Figure S2. Expression of auxin-related genes in response to shade.** Heatmap represents transcript abundance for genes involved in auxin biosynthesis and signaling in the leaves of MM and *hy5-mm2* plants exposed to either W or W + FR for 24 h. See **Dataset S1** for TPM values.

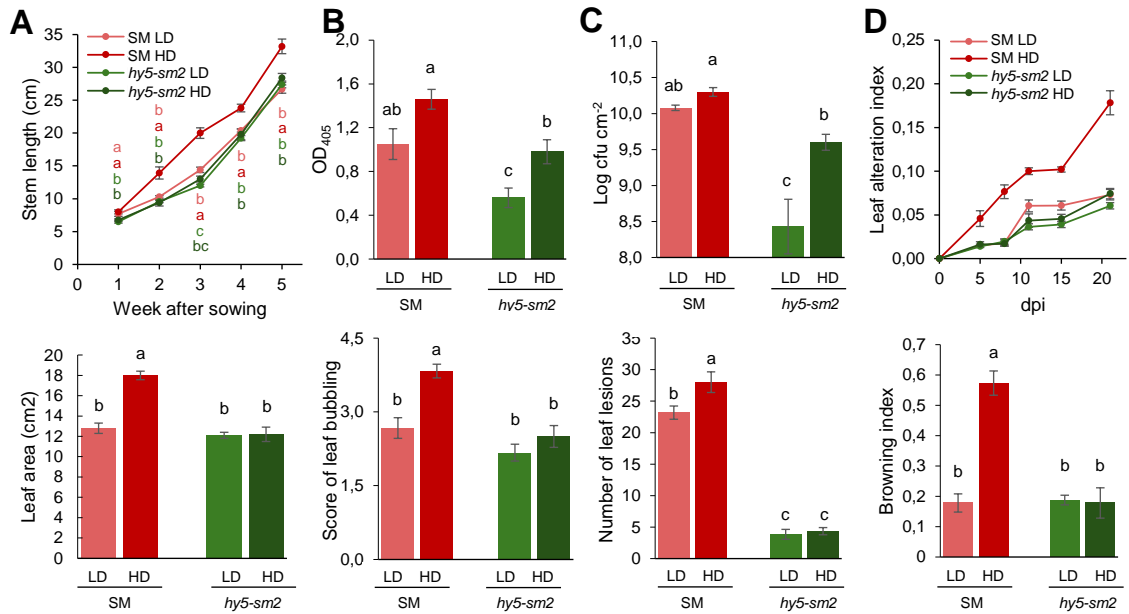

**Figure S3. Same as Figure 6 but with SM and *hy5-sm2* lines.** (A) Stem length and leaf area of tomato wild-type San Marzano (SM) and edited *hy5-sm2* plants grown under low-density (LD) and high-density (HD) conditions in the greenhouse. Stem length was measured weekly and leaf area was measured in 5-week-old plants. (B) Response to *PepMV* infection, evaluated by viral accumulation using an ELISA assay (OD<sub>405</sub>) and by leaf bubbling symptom scores. (C) Response to *Pst* infection, evaluated by bacterial population (log<sub>10</sub> CFU cm<sup>-2</sup>) and by the number of leaf lesions. (D) Response to *Verticillium dahliae* infection, evaluated using the leaf alteration index and browning index parameters. In all plots, dots represent individual data points and error bars represent SE of a minimum of n=5 independent samples (individual plants) per treatment. Statistically significant differences are represented with letters (two-way ANOVA followed by Duncan's multiple range test, P < 0.05).

### SUPPLEMENTAL TABLES

**Table S1. Sequences of cDNA and predicted reading frames in wild type and edited lines.** Allele designation includes the original parental line (MicroTom, mt; San Marzano, sm; or MoneyMaker, mm ). The position of the sgRNA sequence is underlined. Insertions and deletions are marked in red.

|  | Sequence (82-141 cDNA, 28-48 protein) |  |  |  |  |  |  |  |  |  |  |  |  |  |  |  |  |  |  |  |  |  |  |
| --- | --- | --- | --- | --- | --- | --- | --- | --- | --- | --- | --- | --- | --- | --- | --- | --- | --- | --- | --- | --- | --- | --- | --- |
| HY5 | CAT | GAA | CTC | AAA | GAA | GGT | ATG | GAG | AGT | <u>GAT</u> | <u>GAT</u> | GAG | ATC | AGA | AGA | GTG | CCG | GAG | ATG | GGC | GGA |  |  |
|  | H | E | L | K | E | G | M | E | S | D | D | E | I | R | R | V | P | E | M | G | G |  |  |
| hy5-mt4 | C | ATG | AAC | TCA | AAG | AAG | GTA | TGG | AGA | GTG | <u>ATG</u> | <u>ATG</u> | AGA | TCA | GAA | GAG | <u>TG</u> | CCG | GAG | ATG | GGC | GGA |  |
|  | M | N | S | K | K | V | W | R | V | M | M | R | S | E | E | L | P | E | M | G | G |  |  |
| hy5-sm1 | C | ATG | AAC | TCA | AAG | AAG | GTA | TGG | AGA | GTG | <u>ATG</u> | <u>ATG</u> | AGA | TCA | GAA | GAG | <u>CTG</u> | CCG | GAG | ATG | GGC | GGA |  |
|  | M | N | S | K | K | V | W | R | V | M | M | R | S | E | E | L | P | E | M | G | G |  |  |
| hy5-sm.2 | C | ATG | AAC | TCA | AAG | AAG | GTA | TGG | AGA | GTG | <u>ATG</u> | <u>ATG</u> | AGA | TCA | GAA | GAG | <u>ATG</u> | CCG | GAG | ATG | GGC | GGA |  |
|  | M | N | S | K | K | V | W | R | V | M | M | R | S | E | E | M | P | E | M | G | G |  |  |
| hy5-mm2 | C | ATG | AAC | TCA | AAG | AAG | GTA | TGG | AGA | GTG | <u>ATG</u> | <u>ATG</u> | AGA | TCA | GA | A-- | GTG | CCG | GAG | ATG | GGC | GGA |  |
|  | M | N | S | K | K | V | W | R | V | M | M | R | S | E |  | V | P | E | M | G | G |  |  |
| hy5-mt2 | C | ATG | AAC | TCA | AAG | AAG | GTA | TGG | AGA | GTG | <u>ATG</u> | <u>ATG</u> | AGA | TC | --- | --- | A | GTG | CCG | GAG | ATG | GGC | GGA |
|  | M | N | S | K | K | V | W | R | V | M | M | R | S |  |  | V | P | E | M | G | G |  |  |
| hy5-mt1 | C | ATG | AAC | TCA | AAG | AA | G-- | --- | --- | --- | --- | --- | --- | --- | --- | --- | GTG | CCG | GAG | ATG | GGC | GGA |  |
|  | M | N | S | K | K |  |  |  |  |  |  |  |  |  |  |  | V | P | E | M | G | G |  |
| hy5-mm1 | CA | TGA | ACT | CAA | AGA | AGG | TAT | GGA | GAG | TGA | <u>TGA</u> | <u>TGA</u> | GAT | CAG | AAG | A | G-G | CCG | GAG | ATG | GGC | GGA |  |
|  | * |  |  |  |  |  |  |  |  | * | * | * |  |  |  |  |  |  |  | M | G | G |  |

**Table S2. Primers used in this work.**

| Use | Name | Target | Sequence (5'-3') |
| --- | --- | --- | --- |
| sgRNA | sgRNA-F | sgRNA | ATTGTGATGAGATCAGAAGAGTGC |
|  | sgRNA-R | sgRNA | AAACGCACTCTTCTGATCTCATCA |
| Genotyping | Cas9-F | Cas9 | TCCCTCATCAGATCCACCTC |
|  | Cas9-R | Cas9 | CTGAAACCTGAGCCTTCTGG |
|  | SIHY5_CRISPR-F | SIHY5 | GTCCCGCTATTTCTTTTCTG |
|  | SIHY5_CRISPR-R | SIHY5 | GCCCATAACATGTGACAAG |
|  | SIHY5_CRISPR-seq | SIHY5 | CTACGTGGCATGATGTTTAAG |
| Cloning | SIHY5-F_attb1 | SIHY5 | GGGGACAAGTTTGTACAAAAAAGCAGGCTTTATGCAAGAGCAAGCGACGAGTTC |
|  | SIHY5-R_attb2 | SIHY5 | GGGGACCACTTTGTACAAGAAAGCTGGGTTCTTCCTCCCTTCCTGTGCAC |
|  | SICOP1_F_attb1 | SICOP1 | GGGGACAAGTTTGTACAAAAAAGCAGGCTTTATGGTGGAAAGTTCAGTTGGAGG |
|  | SICOP1_R_attb2 | SICOP1 | GGGGACCACTTTGTACAAGAAAGCTGGGTTAGCTGCAAGGACTAACACTT |
|  | SISPA1_F_attb1 | SISPA1 | GGGGACAAGTTTGTACAAAAAAGCAGGCTTTATGGACAAGTCAAAGGAGGAAGC |
|  | SISPA1_R_attb2 | SISPA1 | GGGGACCACTTTGTACAAGAAAGCTGGGTTTACCAAACGTAACAGCTTTA |
|  | OR_F_attb1 | OR | GGGGACAAGTTTGTACAAAAAAGCAGGCTGGATGTCATCTTTGGGTAGGAT |
|  | OR_R_attb2 | OR | GGGGACCACTTTGTACAAGAAAGCTGGGTCATCGAAAGGGTCGATACGAG |
| qPCR | SIHY5_qPCR_F | SIHY5 | GACGAGTTCTATTGCCGCTAG |
|  | SIHY5_qPCR_R | SIHY5 | CCGATACTCCATCTCTTCCAG |

**Table S3. Constructs and cloning details.**

| Use | Construct | Vector | Template | Cloning method |
| --- | --- | --- | --- | --- |
| CRISPR | pEn-SIHY5 | pEn-Chimera C | MM leaf cDNA | *BbsI* / T4 ligase |
|  | pDe-Cas9-SIHY5 | pDE-Cas9 | pEn-SlHY5 | Gateway |
| Cloning | pDONR207-SIHY5 (pMVT7) | pDONR207 | MM leaf cDNA | Gateway |
|  | pDONR207-hy5-mm1 (pMVT8) | pDONR207 | *hy5-mm1* leaf cDNA | Gateway |
|  | pDONR207-hy5-mm2 (pMVT9) | pDONR207 | *hy5-mm2* leaf cDNA | Gateway |
|  | pDONR207-SICOP1 | pDONR207 | MM leaf cDNA | Gateway |
|  | pDONR207-SlSPA1 | pDONR207 | MM leaf cDNA | Gateway |
|  | pDONR207-OR | pDONR207 | Col-0 seedling cDNA | Gateway |
| Immunoblot | p35S-SIHY5-myc | pGWB417 | pDONR207-SlHY5 | Gateway |
|  | p35S-hy5-mm1-myc | pGWB417 | pDONR207-hy5-mm1 | Gateway |
|  | p35S-hy5-mm2-myc | pGWB417 | pDONR207-hy5-mm2 | Gateway |
| Localization | p35S-SIHY5-GFP (pMVT10) | pMDC83 | pDONR207-SlHY5 | Gateway |
|  | p35S-hy5-mm1-GFP (pMVT11) | pMDC83 | pDONR207-hy5-mm1 | Gateway |
|  | p35S-hy5-mm2-GFP (pMVT12) | pMDC83 | pDONR207-hy5-mm2 | Gateway |
| BiFC | YFN-SIHY5 | YFN43-GW | pDONR207-SlHY5 | Gateway |
|  | YFN-hy5-mm2 | YFN43-GW | pDONR207-hy5-mm2 | Gateway |
|  | YFN-SlCOP1 | YFN43-GW | pDONR207-SlCOP1 | Gateway |
|  | YFN-SlSPA1 | YFN43-GW | pDONR207-SlSPA1 | Gateway |
|  | YFN-OR | YFN43-GW | pDONR207-AtOR | Gateway |
|  | YFC-SIHY5 | YFC43-GW | pDONR207-SlHY5 | Gateway |
|  | YFC-hy5-mm2 | YFC43-GW | pDONR207-hy5-mm2 | Gateway |
|  | YFC-SlCOP1 | YFC43-GW | pDONR207-SlCOP1 | Gateway |
|  | YFC-SlSPA1 | YFC43-GW | pDONR207-SlSPA1 | Gateway |
|  | YFC-OR | YFC43-GW | pDONR207-AtOR | Gateway |
